# A dominant *SlTDM1* allele leads to unreduced gametes and polyploid offspring in tomato

**DOI:** 10.64898/2026.11.18.767649

**Authors:** Iris E. Zahn, Freija L.S. Neuteboom Spijker, Maarten P. Wessel, Sofia Ortega, Nicole Jansen, Aleksandr Seleznev, Marleen W. Van der Wiel, Bas Heidemann, Charles J. Underwood

**Affiliations:** Department of Plant & Animal Biology, Radboud Institute for Biological and Environmental Sciences, Radboud University, Nijmegen, the Netherlands

**Keywords:** Meiosis, TDM1, ploidy, cell division, phosphorylation, *mitosis instead of meiosis* (*MiMe*), Tomato, Solanum lycopersicum

## Abstract

Engineering clonal reproduction through seeds typically requires the production of unreduced (2n) clonal gametes. Unreduced clonal gametes can be engineered through the *Mitosis instead of Meiosis* (*MiMe*) system, which so far has been developed in *Arabidopsis*, rice, tomato and *Brassica napus*. However, there is no universal genetic strategy to engineer *MiMe* as the factors regulating the number of meiotic cell divisions show functional divergence. A potential universal factor to abolish the second meiotic cell division is *THREE DIVISIONS MUTANT1* (*TDM1*), which promotes the termination of meiosis in *Arabidopsis* and maize. Here, we developed transgenic tomato plants that express a tomato *TDM1* allele (*SlTDM1-P19L*), which is expected to encode a phosphorylation-defective SlTDM1. We observed that *SlTDM1-P19L* lines produced a mixture of haploid and diploid gametes, which gave rise to up to 28% tetraploid selfing offspring from diploid parents with only mild effects on male fertility. In one line, the *SlTDM1-P19L* transgene led to co-suppression of the endogenous *SlTDM1* gene, resulting in the production of hyper-reduced polyads rather than unreduced dyads or reduced tetrads, mimicking *tdm1* null mutants. Together, our results show that the function of *TDM1* is conserved in tomato, *SlTDM1-P19L* can be effective in producing unreduced gametes, and it is a promising factor to apply in *MiMe* system engineering.

**Significance statement:** Engineering clonal gametes by *Mitosis instead of Meiosis* (*MiMe*) requires abolishing the second meiotic division without compromising fertility, yet universal targets are absent to date. Here, we demonstrate that SlTDM1 regulates meiotic exit in tomato and that expressing a phosphorylation-defective *SlTDM1-P19L* allele produces unreduced gametes and tetraploid offspring, establishing TDM1 as a promising target for establishing *MiMe* in crop species.

## Introduction

Engineering clonal seed production in crops typically requires modifying the meiotic cell cycle (Heidemann et al. 2025). During regular meiosis, two cell divisions result in haploid (1n) spores, giving rise to haploid gametes, thereby ensuring each cycle of sexual reproduction leads to consistent somatic ploidy levels (2n) over generations (Mercier et al. 2015). In contrast, during apomeiosis, a modified meiotic cell division replaces regular meiosis, resulting in clonal, unreduced (2n) spores. The resulting unreduced egg cell can be triggered to undergo parthenogenesis and give rise to clonal offspring in absence of fertilization, a process called gametophytic apomixis (Ozias-Akins and Van Dijk 2007). Clonal offspring, and thus also unreduced clonal gametes, are desired in crop biotechnology to fix the hybrid genotype of plants displaying hybrid vigour.

Unreduced clonal female gametes occur in naturally apomictic species such as in the common dandelion (*Taraxacum officinale*) through diplospory (Asker and Jerling 1992; Koltunow 1993), which is controlled by the dominant *DIPLOSPOROUS* locus (Vijverberg et al. 2010). Unreduced clonal gametes can also be engineered by the *Mitosis instead of Meiosis* (*MiMe*) system (d’Erfurth et al. 2009). *MiMe* involves avoiding three steps to turn meiosis into apomeiosis: meiotic recombination initiation; monopolar orientation and segregation of the sister chromatids at the end of the first meiotic division; and entry into the second meiotic division. So far, *MiMe* has successfully been engineered in *Arabidopsis thaliana* (d’Erfurth et al. 2009), rice (Mieulet et al. 2016), tomato (Wang et al. 2024) and *Brassica napus* (Geng et al. 2026). To achieve apomeiosis, two or three genes are mutated in each species. Clonal gametes have applications in synthetic apomixis and polyploid genome design (Wang et al. 2024; Wang and Underwood 2023), and yet there is currently no universal genetic strategy to engineer *MiMe*.

While the genes involved in the first two steps of *MiMe* are conserved and likely functional to achieve *MiMe* in any plant species of interest (d’Erfurth et al. 2009; Geng et al. 2026; Mieulet et al. 2016; Underwood and Mercier 2022; Wang et al. 2024), there is no universal target gene to avoid entry into the second meiotic division. For this last step, the *MiMe* systems in *Arabidopsis* and rice use a mutation of *OMISSION OF SECOND DIVISION1* (*OSD1*) (d’Erfurth et al. 2009; Mieulet et al. 2016), in tomato *TARDY ASYNCHRONOUS MEIOSIS* (*TAM*)/cyclin *CYCA1;2* is mutated (Wang et al. 2024), and in *Brassica napus* such a mutation is not required (Geng et al. 2026), possibly due to its allopolyploid nature. The *Arabidopsis* and rice *MiMe* plants produce dyads at a high frequency and retain high fertility that is comparable to non-mutant plants (d’Erfurth et al. 2009; Mieulet et al. 2016). The tomato *MiMe* genotype is also highly penetrant (93% tetraploid offspring), but fertility is significantly reduced, probably due to the partial penetrance of *Sltam* (~12% tetraploid offspring in *Sltam* single mutants) (Wang et al. 2024, 2026). As the other two *MiMe* mutations (*Slspo11-1 Slrec8*) are highly penetrant and result in early segregation of the sister chromatids, the *MiMe* genotype is lethal for the meiocyte unless a premature termination of meiosis (due to *Sltam* mutation) occurs to prevent aneuploidy. Also in *Arabidopsis, Attam* is not fully penetrant and *Atosd1* outperforms *Attam* in dyad frequency (Cromer et al. 2012; d’Erfurth et al. 2010). However, mutating *OSD1* is not equally effective for *MiMe* engineering in all species, including tomato where it is single-copy and likely plays a role in mitotic cell division (Di et al. 2022; Pang et al. 2024; Sun et al. 2025; Wang et al. 2024; Zhao et al. 2026). Thus, a more effective mutation than *tam* or *osd1* to generate unreduced gametes is crucial to improve the current tomato *MiMe* system.

Other than *OSD1* and *TAM, THREE DIVISIONS MUTANT1/MALE STERILE5* (*TDM1*/*MS5*) controls the number of meiotic cell divisions in *Arabidopsis* and maize (Bulankova et al. 2010; Cifuentes et al. 2016; Cromer et al. 2012; Glover et al. 1998; Ross et al. 1997; Zhang et al. 2024). TDM1 is a plant-specific tetratricopeptide-domain protein that promotes meiotic exit. In both *Arabidopsis* and maize, TDM1 interacts with the anaphase-promoting complex/cyclosome (APC/C), which is an E3 ubiquitin ligase that marks cell cycle proteins for degradation (Cifuentes et al. 2016; Zhang et al. 2024). Moreover, AtTDM1 functions as a translational repressor at the end of meiosis by sequestering components of the translational machinery in meiotic bodies (M-bodies) by interacting with SUPPRESSOR WITH MORPHOGENETIC EFFECTS ON GENITALIA7 (SMG7) (Bulankova et al. 2010; Cairo et al. 2022). Consequently, *tdm1* meiocytes continuously divide, making them hyper-reduced, aneuploid and sterile (Cifuentes et al. 2016; Zhang et al. 2024).

AtTDM1 activity is regulated by phosphorylation. During meiosis I, AtTDM1 is inactivated by phosphorylation by cyclin-dependent kinase A;1 (CDKA;1)-TAM. The phosphate groups abolish the interaction with the APC/C subunits (Cifuentes et al. 2016) and SMG7 (Cairo et al. 2022). Interestingly, when mutating the AtTDM1 phosphorylation site regulating the interaction with the APC/C subunits (T16-P17), AtTDM1 is active during meiosis I and prematurely terminates meiosis. Thus, in contrast to the *Attdm1* null mutant that produces hyper-reduced polyads by going through multiple rounds of meiosis, *AtTDM1-T16A* and *AtTDM1-P17L* alleles produce unreduced dyads by a premature termination of meiosis (Cifuentes et al. 2016). It is likely that in maize, *ZmTDM1* fulfils a similar role as in *Arabidopsis*, as their gene sequences are conserved and ZmTDM1 also interacts with the APC/C subunits (Zhang et al. 2024).

With the aim to further optimize the tomato *MiMe* system and understand whether *TDM1* allele engineering could be a promising strategy for *MiMe* more generally, here we explored the function of the T18-P19 phosphorylation site of SlTDM1 in tomato. We observed that, like in *Arabidopsis*, the phosphorylation-motif mutants produced unreduced gametes, resulting in up to 28% tetraploid offspring from diploid parents. This finding provides proof-of-principle of the function of SlTDM1 and shows the potential of *TDM1* allele engineering towards unreduced clonal gametes.

## Results

### TDM1 phosphorylation-motif mutants produce pollen of varying sizes

To study the progression of meiosis in TDM1 phosphorylation-motif mutants, we generated two constructs containing *TDM1* with point mutations surrounding a phosphorylation site (Fig. 1a). We generated a variant of *SlTDM1* (*Solyc06g075640*) containing a proline-to-leucine substitution at position 19 (*SlTDM1-P19L*), expressed under the control of its native promoter and 3’ UTR. The other construct contained a variant of *Arabidopsis TDM1* (*AT4G20900*) in which the homologous proline was mutated to a leucine (*AtTDM1-P17L*), which was regulated by the *AtTDM1* promoter and 3’ UTR. This AtTDM1-P17L variant terminates meiosis after the first cell division in *Arabidopsis* (Cifuentes et al. 2016). We hypothesized that this *Arabidopsis* clone can also activate the tomato APC/C and thereby prematurely terminate meiosis. Both binary constructs were stably transformed into the inbred tomato variety Micro-Tom (Scott and Harbaugh 1989).

**Figure 1:**
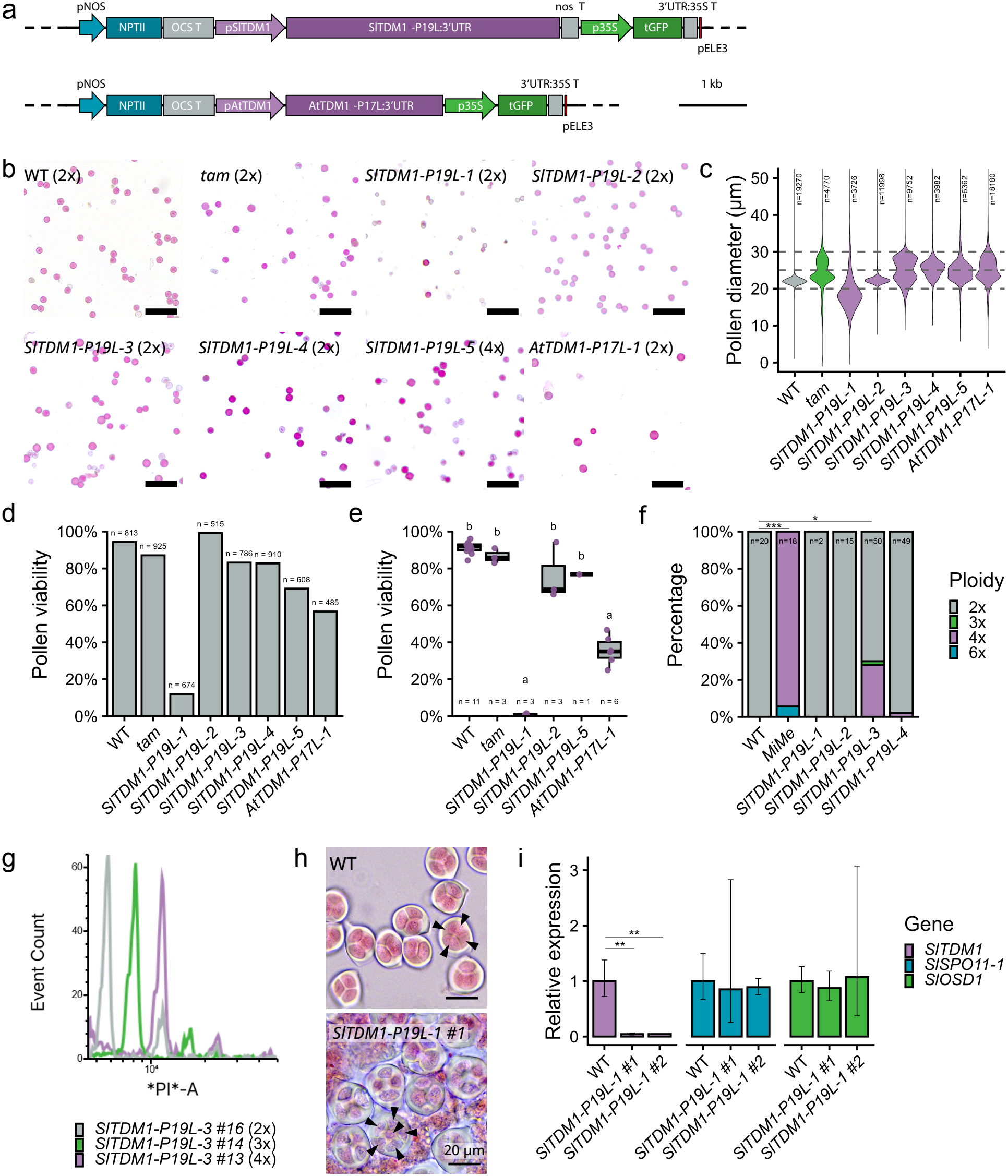
A *SlTDM1-P19L* transgene can trigger unreduced gametes and polyploid offspring in tomato. **(a)** Binary vectors that were generated to introduce a *TDM1* version with a mutated phosphorylation motif. **(b)** Representative images of Alexander staining of pollen from diploid WT (Micro-Tom), diploid *Sltam-1*, five diploid T0 plants (*SlTDM1-P19L-1,-2,-3,-4* and *AtTDM1-P17L-1*) and one tetraploid T0 plant (*SlTDM1-P19L-5*). The scale bars are 100 µm. **(c)** Pollen diameter of diploid WT (Micro-Tom), diploid *Sltam-1*, five diploid T0 plants (*SlTDM1-P19L-1,-2,-3,-4* and *AtTDM1-P17L-1*) and one tetraploid T0 plant (*SlTDM1-P19L-5*) determined by high-throughput image analysis. Pollen between 20-25 µm are classed as haploid and pollen between 25-30 µm are classed as diploid. Sample sizes (n) indicate the number of individual pollen grains measured from a single flower. **(d)** Pollen viability of T0 plants determined by Alexander staining. Sample sizes (n) indicate the number of manually scored pollen. **(e)** Pollen viability of T0 plants determined by impedance flow cytometry. Samples sizes **(f)** indicate the number of analysed flowers. Different letters indicate statistically significant differences. **(g)** Ploidy of T1 plants measured by flow cytometry. * *P*_*adj*_ < 0.05; *** *P*_*adj*_ < 0.001. **(h)** Flow cytometry plot of three T1 *SlTDM1-P19L-3* plants, including one diploid (2x) plant, one triploid (3x) plant and one tetraploid (4x) plant. **(i)** Acetocarmine staining of male meiocytes of WT and *SlTDM1-P19L-1 #1* (T1). Arrows indicate encased microspores. The scale bars are 20 µm. **(j)** Normalized expression of *SlTDM1, SlSPO11-1* and *SlOSD1* in anthers from 3-5 mm flower buds of WT and *SlTDM1-P19L-1* (T1), containing male meiocytes (n=5 for WT and n=2 for *SlTDM1-P19L-1 #1* and *#2*). ** *P* < 0.01. For panels (**b-e**) the lines WT, *tam, SlTDM1-P19L-1* to *-4, AtTDM1-P17L-1* are 2x; *SlTDM1-P19L-5* is 4x. WT refers to Micro-Tom, *tam* refers to *Sltam-1 (*Micro-Tom) (Wang et al., 2024). For (**f**) the parental plant of the *MiMe* offspring was a diploid *Slspo11-1, Slrec8, Sltam* triple mutant (Moneyberg-TMV x Micro-Tom background) (Wang et al., 2024).

We observed that the primary transformants (T0) of both *SlTDM1-P19L* and *AtTDM1-P17L* produced pollen of varying sizes, in contrast to consistently sized pollen of WT plants (Fig. 1b and Figure S1). We identified five independent diploid lines (*SlTDM1-P19L-1* to *-4* and *AtTDM1-P17L-1*) and one tetraploid line (*SlTDM1-P19L-5*) (Figure S2). Image analysis of pollen treated with Alexander’s stain showed a bimodal distribution of smaller haploid (reduced) and larger diploid (unreduced) pollen in the lines *SlTDM1-P19L-3, SlTDM1-P19L-4* and *AtTDM1-P17L-1*, while WT produced only haploid pollen (Fig. 1b, c). Reassuringly the *Sltam-1* mutant (also in Micro-Tom background) produced a mixture of haploid and diploid pollen as expected (Wang et al. 2024), whereas *SlTDM1-P19L-1* produced pollen of low viability which could be consistent with co-suppression of the endogenous *SlTDM1* gene in this line (Fig. 1b, c). In addition, the diploid *SlTDM1-P19L-2* produced viable, haploid pollen like WT, and the viable pollen produced by tetraploid *SlTDM1-P19L-5* was mostly diploid, which could be consistent with low or no transgene activity in these lines. These data are consistent with skipping of the second meiotic division, at partial penetrance, in the male meiocytes of the *SlTDM1-P19L-3, SlTDM1-P19L-4* and *AtTDM1-P17L-1* phosphorylation-motif mutant lines, resulting in a mixture of both haploid (reduced) and diploid (unreduced) pollen.

As modifying meiosis can affect fertility and *tdm1* null mutants are sterile (Glover et al. 1998; Ross et al. 1997; Zhang et al. 2024), we next quantified the viability of the pollen of the TDM1 phosphorylation-motif mutants. We quantified the pollen viability based on the Alexander staining (Fig. 1d) and confirmed these measurements for a selection of lines using impedance flow cytometry (Fig. 1e). In the lines showing variation in pollen sizes (*SlTDM1-P19L-3, SlTDM1-P19L-4* and *AtTDM1-P17L-1*), pollen viability was mildly reduced and ranged between 57% and 83%, compared to 94% in WT (Fig. 1d). Quantification of pollen viability of *SlTDM1-P19L-1* using both methods indicated highly reduced viable pollen, indicating a meiotic defect may occur in this line (Fig. 1d, e).

### Unreduced gametes in *SlTDM1-P19L* give rise to polyploid progeny

To investigate whether the larger pollen of the diploid parents are indeed diploid and give rise to viable progeny, we analysed the ploidy levels of the next generation (T1) (Fig. 1f, g and Table 1). Interestingly, 28% of the *SlTDM1-P19L-3* progeny was tetraploid, which would be consistent with roughly 50% unreduced female gametes and 50% unreduced male gametes in the parent. Moreover, we identified a triploid plant in this line (Fig. 1g), which shows that both haploid and diploid gametes were produced in the same flower and the seed escaped the triploid block (Köhler et al. 2010). In *SlTDM1-P19L-4*, 2% of the progeny was tetraploid. Consistent with the absence of unreduced pollen in *SlTDM1-P19L-1* and *-2* T0 plants, we did not identify any tetraploid T1 progeny in the *SlTDM1-P19L-1* and *-2* lines. We additionally assessed the ploidy of the offspring of a control diploid *Slspo11-1 Slrec8 Sltam* mutant in the Moneyberg-TMV x Micro-Tom F1 hybrid background which we hereafter refer to as *MiMe* (Wang et al. 2024). We found that 94% of the MiMe offspring were tetraploid as only unreduced gametes are viable in the *MiMe* genotype, and is consistent with the previous findings (Wang et al. 2024). Together, this shows that SlTDM1-P19L can lead to unreduced, functional male and female gametes in tomato plants which when they fuse give rise to tetraploid offspring.

**Table 1:**
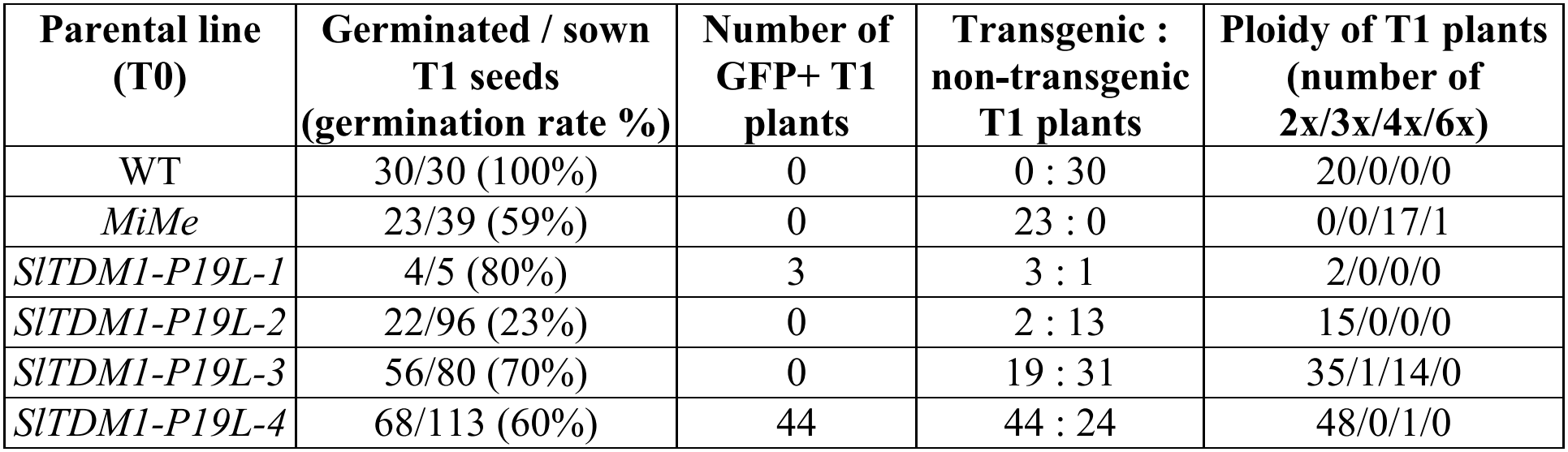
Germination rate, genotyping and ploidy of T1 plants.

### *SlTDM1* silencing leads to hyper-reduced polyads

When germinating the T1 seeds, we noticed that the GFP of the transgene was silenced in the lines *SlTDM1-P19L-2* and *-3* (Table 1 and Figure S3). We also observed that none of the T1 lines produced diploid pollen (Figure S4), suggesting that the *SlTDM1-P19L* transgene was silenced. In the line *SlTDM1-P19L-1*, which had low pollen viability in the T0 and T1 generations, we observed the production of hyper-reduced polyads in T1 plants, in contrast to the reduced tetrads that occur in WT plants (Fig. 1h). These polyads mimic a *tdm1* loss-of-function phenotype in which meiosis continues after the regular two cell divisions (Bulankova et al. 2010; Cifuentes et al. 2016; Cromer et al. 2012; Glover et al. 1998; Ross et al. 1997; Zhang et al. 2024). In line with the polyads, the pollen of *SlTDM1-P19L-1* are mostly smaller than 20 µm and unviable (Figure S4). These observations suggest that silencing of the *SlTDM1-P19L* transgene led to co-suppression of the endogenous *SlTDM1* gene in *SlTDM1-P19L-1*.

To explain the low pollen viability and polyad phenotype in *SlTDM1-P19L-1*, we verified the *SlTDM1* expression levels in anthers undergoing meiosis by qPCR (Fig. 1i). In two *SlTDM1-P19L-1* T1 plants, *SlTDM1* expression was almost completely abolished (Fig. 1i). Two other meiosis genes, *SlSPO11-1*, required for DNA double-strand break formation at the start of meiosis (Grelon et al. 2001), and *SlOSD1*, marking the transition from meiosis I to meiosis II (Cromer et al. 2012), were expressed at WT levels in the mutants, indicating meiotic cells were present in the tested anthers (Fig. 1i). These data support that the *SlTDM1-P19L* transgene in line *SlTDM1-P19L-1* led to the co-suppression of the endogenous *SlTDM1* gene. The loss of *SlTDM1* expression explains the failure to terminate meiosis and thereby led to the production of polyads and reduced fertility.

## Discussion

A crucial step of engineering gametophytic apomixis is the production of unreduced clonal gametes. Here, we explored the possibility to express *TDM1* variants that are expected to produce phosphorylation-defective TDM1 proteins (Brown et al. 1999) and ultimately produce unreduced gametes in tomato. Two diploid independent lines produced tetraploid offspring (Table 1 and Fig. 1f), supporting that SlTDM1 P19 phosphorylation is likely required for the interaction with the APC/C and progression of meiosis in tomato, like in *Arabidopsis* (Cifuentes et al. 2016). The conserved function of TDM1 in promoting meiotic exit is further supported by the production of hyper-reduced meiotic polyads in another line, in which *SlTDM1* expression was abolished (Fig. 1h and Fig. 1i). This strongly resembles the phenotype of *tdm1* null mutants in *Arabidopsis* and maize (Glover et al. 1998; Ross et al. 1997; Zhang et al. 2024). Thus, SlTDM1 appears to have a conserved function in regulating the number of meiotic cell divisions in solanaceous plants.

Although two lines produced unreduced offspring, we encountered transgene silencing and lost the desired effect on meiosis in subsequent generations. We observed that expression of the *p35S::tGFP* gene of the multigene construct was not stable, which could have led to silencing of the linked *SlTDM1-P19L* gene (Roca Paixao and Déléris 2024; Vaucheret et al. 1998). Gene silencing occurs more frequently in transgenes than endogenous genes (Rajeevkumar et al. 2015; Zhang et al. 2015), but can also lead to co-suppression of their homologous endogenous genes, which is famously implemented by trans-acting small RNA molecules (De Felippes and Waterhouse 2020; Hamilton and Baulcombe 1999; Napoli et al. 1990; Van Der Krol et al. 1990). We observed transgene silencing from the T1 generation onwards and in one line this led to the co-suppression of the endogenous *SlTDM1* gene. Therefore, to avoid silencing of *SlTDM1* and apply it in a *MiMe* system, a more stable genome engineering approach should be used. Base editing (Hillary and Ceasar 2024) and prime editing (Van Vu et al. 2026) are becoming more efficient in plants and would be promising strategies.

*TDM1* is a promising factor to apply in a *MiMe* system in species in which *OSD1* cannot be used, which probably includes most species outside the Brassicaceae and Poaceae (Underwood and Mercier 2022). In contrast to *OSD1, TDM1* appears to have a conserved function in monocots and dicots (Bulankova et al. 2010; Cifuentes et al. 2016; Cromer et al. 2012; Glover et al. 1998; Ross et al. 1997; Zhang et al. 2024). *TDM1* is also meiosis specific, as we and others (Cifuentes et al. 2016) did not observe any non-meiotic effects in the TDM1 phosphorylation-motif mutants. Moreover, our results also suggest that in tomato, *SlTDM1-P19L* can be more effective in producing unreduced offspring than *Sltam-1* (Wang et al. 2024), possibly because of the downstream position of TDM1 in the regulation of meiotic exit. Notably the *SlTDM1-P19L-3* line with the highest penetrance (28% tetraploid offspring) only showed a mild decrease in pollen viability (83% compared to 94% in WT). Overall, our results demonstrate that *SlTDM1* has the potential to improve the penetrance and fertility of the current tomato *MiMe* system.

Manipulating meiosis remains a sensitive genetic bottleneck. To obtain unreduced gametes, the first meiotic cell division should proceed as normal and only the second division should be abolished, requiring a very specific effect on the meiotic cell cycle. Furthermore, many core cell-cycle regulators function in both meiosis and mitosis (Wijnker and Schnittger 2013), thereby limiting the factors for meiosis-specific cell cycle engineering. Even among the meiosis-specific regulators, several genes regulate, or are studied in, the male or female meiosis only and are therefore less useful to apply in a *MiMe* system (e.g. (d’Erfurth et al. 2008; Erilova et al. 2009; Ravi et al. 2008; Sofroni et al. 2020)). Future identification of meiosis-specific cyclins and CDKs downstream of the APC/C may offer a more effective strategy for engineering a single meiotic cell division.

## Materials and Methods

### Plant materials and growing conditions

Tomato (*Solanum lycopersicum*) cv. Micro-Tom was used in this study (Scott and Harbaugh 1989). Seeds were treated with 150 mg/L GA_3_ for 20 hours, sterilized using 1% bleach and germinated on 0.7% phyto agar. Seedlings were grown *in vitro* on ½ MS medium (2.15 g/L MS, 50 mg/L myo-inositol, 15 g/L sucrose, 125 mg/L carbenicillin, 7 g/L phyto agar, pH 5.9) at 25 ºC. Seven weeks after germination, plants were transferred to a greenhouse and grown on soil. Moneyberg-TMV x Micro-Tom F1 hybrid *MiMe* (*Slspo11-1 Slrec8 Sltam*) plants and Micro-Tom *tam-1* mutants were reported by Wang et al. (2024).

### Cloning and plant transformations

The *SlTDM1-P19L* and *AtTDM1-P17L* constructs were cloned using the standard Golden Gate cloning strategy (Engler et al. 2014). The *SlTDM1* promoter (1022 bp), gene sequence and 3’ UTR (822 bp) were amplified separately from gDNA by PCR. The proline-to-leucine point mutation was introduced by overlap extension PCR. The fragments were cloned into pICH41295, pAGM1287 and pAGM1301, respectively, and combined into pICH47742. The *AtTDM1-P17L* sequence was amplified from the original plasmid constructed by Cifuentes et al. (2016), domesticated, and cloned into pICH47742. The two final constructs were assembled in pICSL4723 and contained *pNOS::NPTII:OCST*, either *TDM1* sequence, *p35S::tGFP:35ST* and *pELE3*. Primers are listed in Table S1.

Stable tomato transformations with the binary level 2 vectors *SlTDM1-P19L* and *AtTDM1-P17L* were performed according to Van Roekel et al. (1993) with minor alterations. We used the *Agrobacterium tumefaciens* strain GV3101, 2 mg/L zeatin instead of zeatin riboside, and 0.05 mg/L 2,4-D instead of IAA in medium B. Explants were cultured at 25 ºC with a 16h/8h light/dark cycle and a light intensity of 100 µE at plate level. Diploid, transgenic T0 plants were selected by flow cytometry and PCR. Genotyping primers are listed in Table S1.

### Ploidy analysis

Ploidy levels were determined by flow cytometry using a BD FACSMelody Cell Sorter. Nuclei were extracted from a young tomato leaf by chopping the leaf with a razor blade in 500 µL Galbraith’s buffer (45 mM MgCl2, 30 mM sodium citrate, 20 mM MOPS, 0.1% (v/v) Triton X-100, pH 7.0) (Galbraith et al. 1983). The extract was filtered through a 50 µm CellTrics filter. Nuclei were stained with propidium iodide or DAPI. Ploidy levels are based on measurements of ~1,000 nuclei. Data were visualised in Floreada.io. Ploidy distributions of diploid (2n) vs. non-diploid (3n, 4n or 6n) were compared between each transgenic line and the WT using one-sided Fisher’s exact tests. P-values from the five pairwise comparisons with WT were corrected for multiple testing using the Holm method. All statistical analysis were performed in R.

### Microscopy of male meiocytes and pollen

To stain male meiocytes, flower buds of 3-5 mm in length were fixed in 3:1 ethanol : glacial acetic acid at 4 ºC. Meiocytes were released by crushing a single anther between the microscope slide and cover slip in 6 µL of 0.75% acetocarmine in 45% acetic acid. Flowers for pollen staining were used directly or fixed in Carnoy’s fixative (6:3:1 ethanol : chloroform : glacial acetic acid) at 4 ºC. The anther cone was cut transversely and pollen were extracted by immersing a slice of the anther cone in 8 µL Alexander staining (9.5% ethanol, 0.1% malachite green in 95% ethanol, 25% glycerol, 0.05% acid fuchsin, 0.005% orange G, 4% glacial acetic acid) on a microscope slide and tapping the sample. Meiocytes and pollen were imaged using a Leica DMC4500.

### Image analysis for pollen size estimation

Individual pollen grains were segmented from microscope images using Cellpose (Stringer et al. 2021). ImageJ with FIJI plugins was used to do object measurements. The Feret diameter of the object was used as an estimation of the pollen grain diameter. The analysis pipeline described by Frencken and colleagues was used to detect pollen sizes (Frencken et al., 2026). Pollen from a single flower were analysed per line.

### Pollen viability assays

Pollen viability was determined by Alexander staining (staining only viable pollen) and impedance flow cytometry (based on electric conductivity of live and dead cells) (Heidmann et al. 2016). The Alexander staining was performed as described above. Per flower, 485-910 pollen were manually scored for viability in a selected region of the microscopy image. For the impedance flow cytometry an Ampha Z32 with a D-chip and AF6 buffer was used according to the manufacturer’s recommendations for tomato pollen. Per measurement, ~1,000 pollen of a single flower at anthesis were analysed. The mean pollen viability of every measurement was used as biological replicate. Statistical differences were calculated using a Kruskal-Wallis test followed by a Dunn’s post-hoc test with Holm correction for pairwise comparisons.

### Expression analysis by qPCR

RNA was extracted from anthers of 3-5 mm flower buds using TRIzol (Invitrogen) according to the manufacturer’s instructions. Each biological replicate contained a pool of 3 anther cones of varying sizes from a single plant. DNA was removed from the samples using the TURBO DNA-free kit (ThermoFisher Scientific). cDNA was synthesized from 400 ng RNA using iScript (BioRad). Real-time qPCR was performed with iQ SYBR Green Supermix (BioRad) in a CFX Connect Real-Time machine (BioRad) with *ACTIN* and *GAPDH* as reference genes. Primers are listed in Table S1. Significant differences were calculated using a two-tailed homoscedastic t-test.

## Acknowledgements

Micro-Tom seeds were originally sourced from the Tomato Growers Supply Company (USA). The original *pAtTDM1::AtTDM1-P17L* construct (Cifuentes et al. 2016) was kindly shared by Raphael Mercier (MPI-PZ, Cologne, Germany). We thank Yazhong Wang who generated the *MiMe* and *tam-1* mutants. We thank Max Frencken and Daan van den Brink for help with the pollen image analysis, and Tobias de Werk and Jan Zethof for assistance with RNA extractions and qPCR. We also thank the Radboud University greenhouse personnel (Koos Janssen, Walter Hendrickx, Peter Arns and Dorine Schonhoven-van Eeuwijk) for the plant care. This work was supported by the Radboud University executive board and the European Research Council (ERC starting grant 101076355, “AsexualEmbryo” to C.J.U.).

## References

Asker S and Jerling L. Apomixis in Plants (CRC Press). 1992.

Brown NR, Noble MEM, Endicott JA, and Johnson LN. The structural basis for specificity of substrate and recruitment peptides for cyclin-dependent kinases. Nat Cell Biol. 1999:1(7):438–443. 10.1038/15674

Bulankova P, Riehs-Kearnan N, Nowack MK, Schnittger A, and Riha K. Meiotic Progression in Arabidopsis Is Governed by Complex Regulatory Interactions between SMG7, TDM1, and the Meiosis I–Specific Cyclin TAM. Plant Cell. 2010:22(11):3791–3803. 10.1105/tpc.110.078378

Cairo A, Vargova A, Shukla N, Capitao C, Mikulkova P, Valuchova S, Pecinkova J, Bulankova P, and Riha K. Meiotic exit in Arabidopsis is driven by P-body–mediated inhibition of translation. Science. 2022:377(6606):629–634. 10.1126/science.abo0904

Cifuentes M, Jolivet S, Cromer L, Harashima H, Bulankova P, Renne C, Crismani W, Nomura Y, Nakagami H, Sugimoto K, et al. TDM1 Regulation Determines the Number of Meiotic Divisions. PLoS Genet. 2016:12(2):e1005856. 10.1371/journal.pgen.1005856

Cromer L, Heyman J, Touati S, Harashima H, Araou E, Girard C, Horlow C, Wassmann K, Schnittger A, De Veylder L, et al. OSD1 Promotes Meiotic Progression via APC/C Inhibition and Forms a Regulatory Network with TDM and CYCA1;2/TAM. PLoS Genet. 2012:8(7):e1002865. 10.1371/journal.pgen.1002865

De Felippes FF and Waterhouse PM. The Whys and Wherefores of Transitivity in Plants. Front Plant Sci. 2020:12(1):579376. 10.3389/fpls.2020.579376

Di S, Zhang P, Zhang J, Liu G, Wang G, Shi Q, Ma F, and Bao Z. Tomato UVI4 homologue modulates cell expansion to participate heat-stimulated hypocotyl elongation. Environmental and Experimental Botany. 2022:201:104963. 10.1016/j.envexpbot.2022.104963

Engler C, Youles M, Gruetzner R, Ehnert T-M, Werner S, Jones JDG, Patron NJ, and Marillonnet S. A Golden Gate Modular Cloning Toolbox for Plants. ACS Synth Biol. 2014:3(11):839–843. 10.1021/sb4001504

d’Erfurth I, Cromer L, Jolivet S, Girard C, Horlow C, Sun Y, To JPC, Berchowitz LE, Copenhaver GP, and Mercier R. The CYCLIN-A CYCA1;2/TAM Is Required for the Meiosis I to Meiosis II Transition and Cooperates with OSD1 for the Prophase to First Meiotic Division Transition. PLoS Genet. 2010:6(6):e1000989. 10.1371/journal.pgen.1000989

d’Erfurth I, Jolivet S, Froger N, Catrice O, Novatchkova M, and Mercier R. Turning Meiosis into Mitosis. PLoS Biol. 2009:7(6):e1000124. 10.1371/journal.pbio.1000124

d’Erfurth I, Jolivet S, Froger N, Catrice O, Novatchkova M, Simon M, Jenczewski E, and Mercier R. Mutations in AtPS1 (Arabidopsis thaliana Parallel Spindle 1) Lead to the Production of Diploid Pollen Grains. PLoS Genet. 2008:4(11):e1000274. 10.1371/journal.pgen.1000274

Erilova A, Brownfield L, Exner V, Rosa M, Twell D, Scheid OM, Hennig L, and Köhler C. Imprinting of the Polycomb Group Gene MEDEA Serves as a Ploidy Sensor in Arabidopsis. PLoS Genet. 2009:5(9):e1000663. 10.1371/journal.pgen.1000663

Frencken M, van den Brink DM, van den Hurk J, van der Krabben L, Serin EAR, and Rieu, I.. Genome-wide association study in Arabidopsis reveals IAN genes regulate pollen heatwave tolerance. Research Square PREPRINT. 2026. 10.21203/rs.3.rs-11319537/v1

Galbraith DW, Harkins KR, Maddox JM, Ayres NM, Sharma DP, and Firoozabady E. Rapid Flow Cytometric Analysis of the Cell Cycle in Intact Plant Tissues. Science. 1983:220(4601):1049–1051. 10.1126/science.220.4601.1049

Geng M, Chu L, Guo Z, Zhang Y, Zhuang J, Xu G, Yuan X, Cai B, and Yang C. Establishing a 2-gene MiMe to produce clonal gametes in allotetraploid Brassica napus. Plant Cell. 2026:38(5):koag145. 10.1093/plcell/koag145

Glover J, Grelon M, Craig S, Chaudhury A, and Dennis E. Cloning and characterization of MS5 from Arabidopsis : a gene critical in male meiosis. Plant J. 1998:15(3):345–356. 10.1046/j.1365-313X.1998.00216.x

Grelon M, Vezon D, Gendrot G, and Pelletier G. AtSPO11-1 is necessary for efficient meiotic recombination in plants. EMBO J. 2001:20(3):589–600. 10.1093/emboj/20.3.589

Hamilton AJ and Baulcombe DC. A Species of Small Antisense RNA in Posttranscriptional Gene Silencing in Plants. Science. 1999:286(5441):950–952. 10.1126/science.286.5441.950

Heidemann B, Primetis E, Zahn IE, and Underwood CJ. To infinity and beyond: recent progress, bottlenecks, and potential of clonal seeds by apomixis. Plant J. 2025:121(4):e70054. 10.1111/tpj.70054

Heidmann I, Schade-Kampmann G, Lambalk J, Ottiger M, and Di Berardino M. Impedance Flow Cytometry: A Novel Technique in Pollen Analysis. PLoS ONE. 2016:11(11):e0165531. 10.1371/journal.pone.0165531

Hillary VE and Ceasar SA. CRISPR/Cas system-mediated base editing in crops: recent developments and future prospects. Plant Cell Rep. 2024:43(11):271. 10.1007/s00299-024-03346-0

Köhler C, Mittelsten Scheid O, and Erilova A. The impact of the triploid block on the origin and evolution of polyploid plants. Trends Genet. 2010:26(3):142–148. 10.1016/j.tig.2009.12.006

Koltunow AM. Apomixis: Embryo Sacs and Embryos Formed without Meiosis or Fertilization in Ovules. Plant Cell. 1993:5(10):1425–1437. 10.1105/tpc.5.10.1425

Mercier R, Mézard C, Jenczewski E, Macaisne N, and Grelon M. The Molecular Biology of Meiosis in Plants. Annu Rev Plant Biol. 2015:66(1):297–327. 10.1146/annurev-arplant-050213-035923

Mieulet D, Jolivet S, Rivard M, Cromer L, Vernet A, Mayonove P, Pereira L, Droc G, Courtois B, Guiderdoni E, et al. Turning rice meiosis into mitosis. Cell Res. 2016:26(11):1242–1254. 10.1038/cr.2016.117

Napoli C, Lemieux C, and Jorgensen R. Introduction of a Chimeric Chalcone Synthase Gene into Petunia Results in Reversible Co-Suppression of Homologous Genes in trans. Plant Cell. 1990:2(4):279–289. 10.1105/tpc.2.4.279

Ozias-Akins P and Van Dijk PJ. Mendelian Genetics of Apomixis in Plants. Annu Rev Genet. 2007:41(1):509–537. 10.1146/annurev.genet.40.110405.090511

Pang W, He W, Liang J, Wang Q, Hou S, Luo X, Li J, Wang J, Tian S, and Yuan L. Disruption of ClOSD1 leads to both somatic and gametic ploidy doubling in watermelon. Horticulture Research. 2024:uhae288. 10.1093/hr/uhae288

Rajeevkumar S, Anunanthini P, and Sathishkumar R. Epigenetic silencing in transgenic plants. Front Plant Sci. 2015:6(693). 10.3389/fpls.2015.00693

Ravi M, Marimuthu MPA, and Siddiqi I. Gamete formation without meiosis in Arabidopsis. Nature. 2008:451(7182):1121–1124. 10.1038/nature06557

Roca Paixao JF and Déléris A. Epigenetic control of T-DNA during transgenesis and pathogenesis. Plant Physiol. 2024:197(1):kiae583. 10.1093/plphys/kiae583

Ross KJ, Fransz P, Armstrong SJ, Vizir I, Mulligan B, Franklin FCH, and Jones GH. Cytological characterization of four meiotic mutants of Arabidopsis isolated from T-DNA-transformed lines. Chromosome Res. 1997:5(8):551–559. 10.1023/A:1018497804129

Scott JW and Harbaugh BK. Micro-Tom. A miniature dwarf tomato. Agricultural Experiment Station, Institute of Food and Agricultural Sciences, University of Florida. 1989:370.

Sofroni K, Takatsuka H, Yang C, Dissmeyer N, Komaki S, Hamamura Y, Böttger L, Umeda M, and Schnittger A. CDKD-dependent activation of CDKA;1 controls microtubule dynamics and cytokinesis during meiosis. J Cell Biol. 2020:219(8):e201907016. 10.1083/jcb.201907016

Stringer C, Wang T, Michaelos M, and Pachitariu M. Cellpose: a generalist algorithm for cellular segmentation. Nat Methods. 2021:18(1):100–106. 10.1038/s41592-020-01018-x

Sun M, Lu J, Wu L, Wang B, Han J, Li Y, Yang H, Xie Q, Wang Z, Yang L, et al. StOSD1/UVI4 is crucial for accurate mitotic progression and the maintenance of leaf morphology in potato. BMC Plant Biol. 2025:25(1):1284. 10.1186/s12870-025-07318-8

Underwood CJ and Mercier R. Engineering Apomixis: Clonal Seeds Approaching the Fields. Annu Rev Plant Biol. 2022:73(1):201–225. 10.1146/annurev-arplant-102720-013958

Van Der Krol AR, Mur LA, Beld M, Mol JN, and Stuitje AR. Flavonoid genes in petunia: addition of a limited number of gene copies may lead to a suppression of gene expression. Plant Cell. 1990:2(4):291–299. 10.1105/tpc.2.4.291

Van Roekel JSC, Damm B, Melchers LS, and Hoekema A. Factors influencing transformation frequency of tomato (Lycopersicon esculentum). Plant Cell Rep. 1993:12(11):644–647. 10.1007/BF00232816

Van Vu T, Thi Nguyen N, Kim J, Hoai Nguyen T, and Kim J-Y. Development of an ultra-efficient prime editing system in tomato. Nat Commun. 2026:17(1):95. 10.1038/s41467-025-67874-3

Vaucheret H, Béclin C, Elmayan T, Feuerbach F, Godon C, Morel J, Mourrain P, Palauqui J, and Vernhettes S. Transgene-induced gene silencing in plants. Plant J. 1998:16(6):651–659. 10.1046/j.1365-313x.1998.00337.x

Vijverberg K, Milanovic-Ivanovic S, Bakx-Schotman T, and Van Dijk PJ. Genetic fine-mapping of DIPLOSPOROUS in Taraxacum (dandelion; Asteraceae) indicates a duplicated DIP-gene. BMC Plant Biol. 2010:10(1):154. 10.1186/1471-2229-10-154

Wang Y, Fuentes RR, Rengs WMJ van, Effgen S, Zaidan MWAM, Franzen R, Susanto T, Fernandes JB, Mercier R, and Underwood CJ. Harnessing clonal gametes in hybrid crops to engineer polyploid genomes. Nature Genetics. 2024. 10.1038/s41588-024-01750-6

Wang Y and Underwood CJ. Apomixis. Curr Biol. 2023:33(8):R293–R295. 10.1016/j.cub.2023.01.051

Wang Y, Zahn IE, Heidemann B, Fuentes RR, and Underwood CJ. Synthetic Apomeiosis in Tomato via the Mutagenesis of SlSPO11-1, SlREC8, and SlTAM to Engineer Mitosis Instead of Meiosis. In. Apomixis, T Radoeva and PJ Van Dijk, eds, Methods in Molecular Biology. (Springer US: New York, NY), pp. 255–273. 10.1007/978-1-0716-5001-1_17

Wijnker E and Schnittger A. Control of the meiotic cell division program in plants. Plant Reprod. 2013:26(3):143–158. 10.1007/s00497-013-0223-x

Zhang T, Zhao S, and He Y. ZmTDM1 encodes a tetratricopeptide repeat domain protein and is required for meiotic exit in maize. Plant J. 2024:117(5):1517–1527. 10.1111/tpj.16579

Zhang X, Zhu Y, Liu X, Hong X, Xu Y, Zhu P, Shen Y, Wu H, Ji Y, Wen X, et al. Suppression of endogenous gene silencing by bidirectional cytoplasmic RNA decay in Arabidopsis. Science. 2015:348(6230):120–123. 10.1126/science.aaa2618

Zhao T, Zhang Y, Wu J, Hussain I, Li W, Ping F, Liu K, Pan C, and Yu X. Development of a low-penetrance mitosis instead of meiosis system in tomato. Mol Horticulture. 2026:6(1):38. 10.1186/s43897-025-00207-6

