## Supplementary information for "A dominant *SlTDM1* allele leads to unreduced gametes and polyploid offspring in tomato"

**Supplementary data**

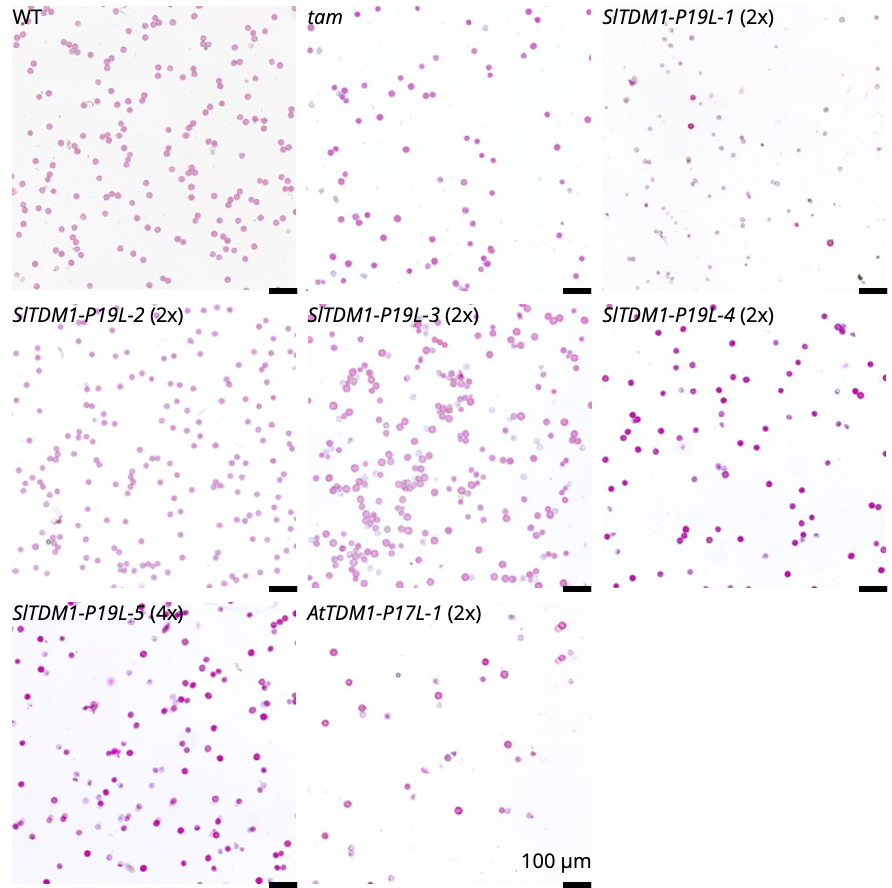

**Figure. S1: Alexander staining of pollen from WT, *tam*, diploid T0 plants (*SlTDM1-P19L-1* to *-4* and *AtTDM1-P17L-1*) and one tetraploid T0 plant (*SlTDM1-P19L-5*).**

The scale bars are 100 µm. WT refers to Micro-Tom and *tam* refers to *Sltam-1 (*Micro-Tom) (Wang et al., 2024).

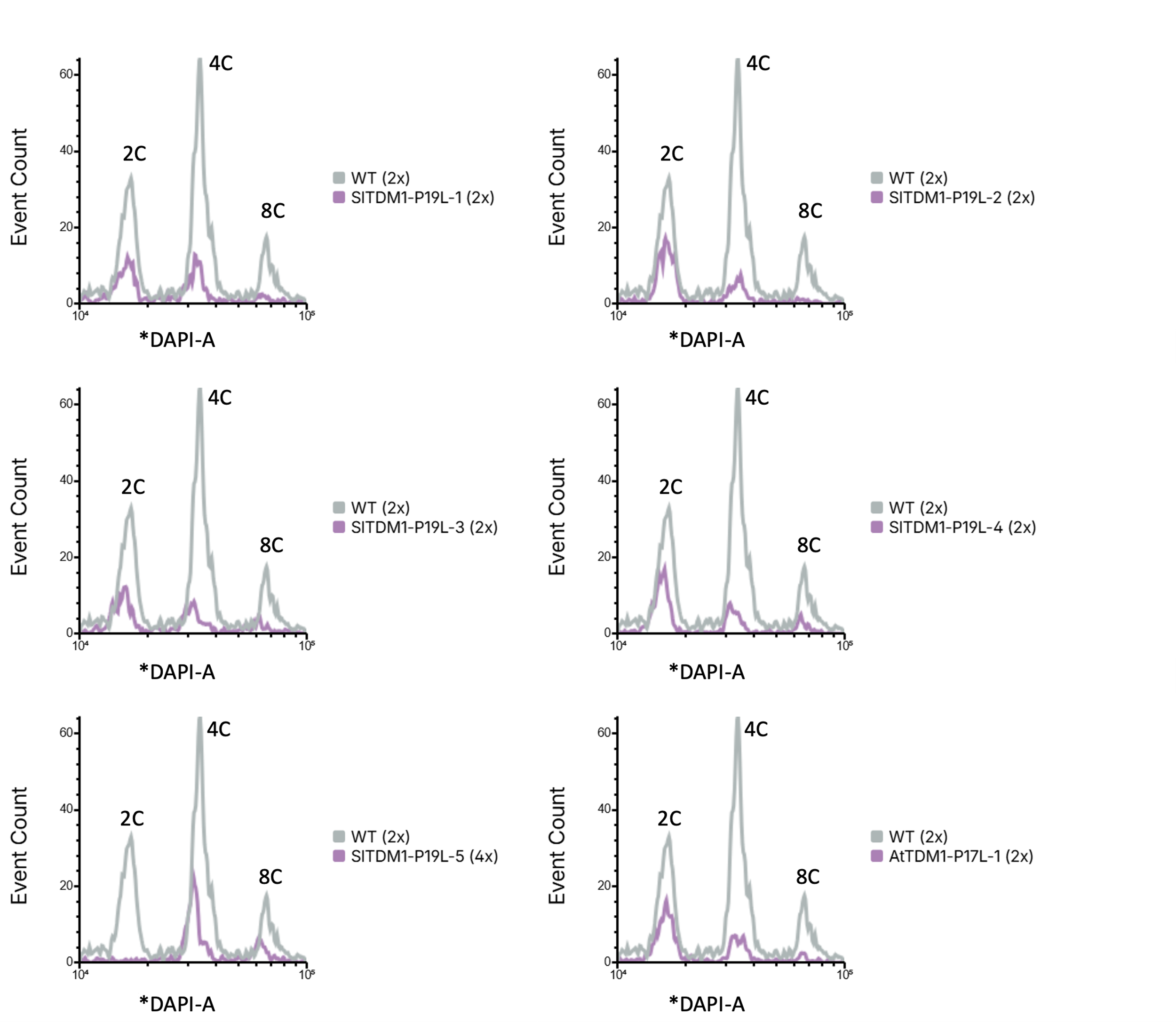

**Figure S2: Flow cytometry data of T0 plants *SlTDM1-P19L-1*, *SlTDM1-P19L-2* *SlTDM1-P19L-3*, *SlTDM1-P19L-4*, *SlTDM1-P19L-5* and *AtTDM1-P17L-1*.**

The flow cytometry profiles of the respective transgenic lines are overlaid with a single diploid WT control. WT refers to Micro-Tom.

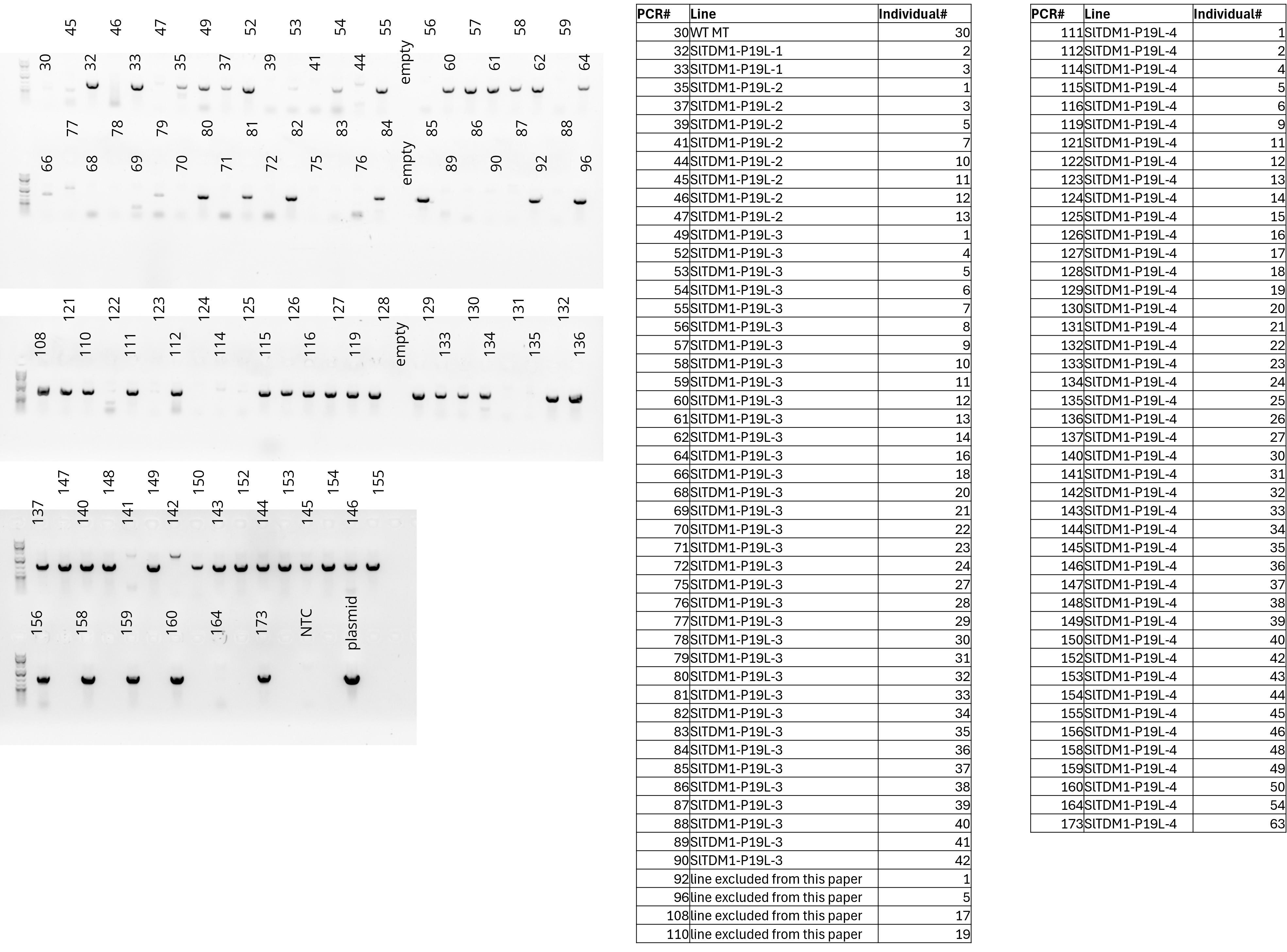

**Fig. S3: Genotyping of T1 plants for the presence of the *SlTDM1* construct**

The numbers on the gel image correspond to the PCR# in the table. The line refers to the T0 parent. NTC: non-template control.

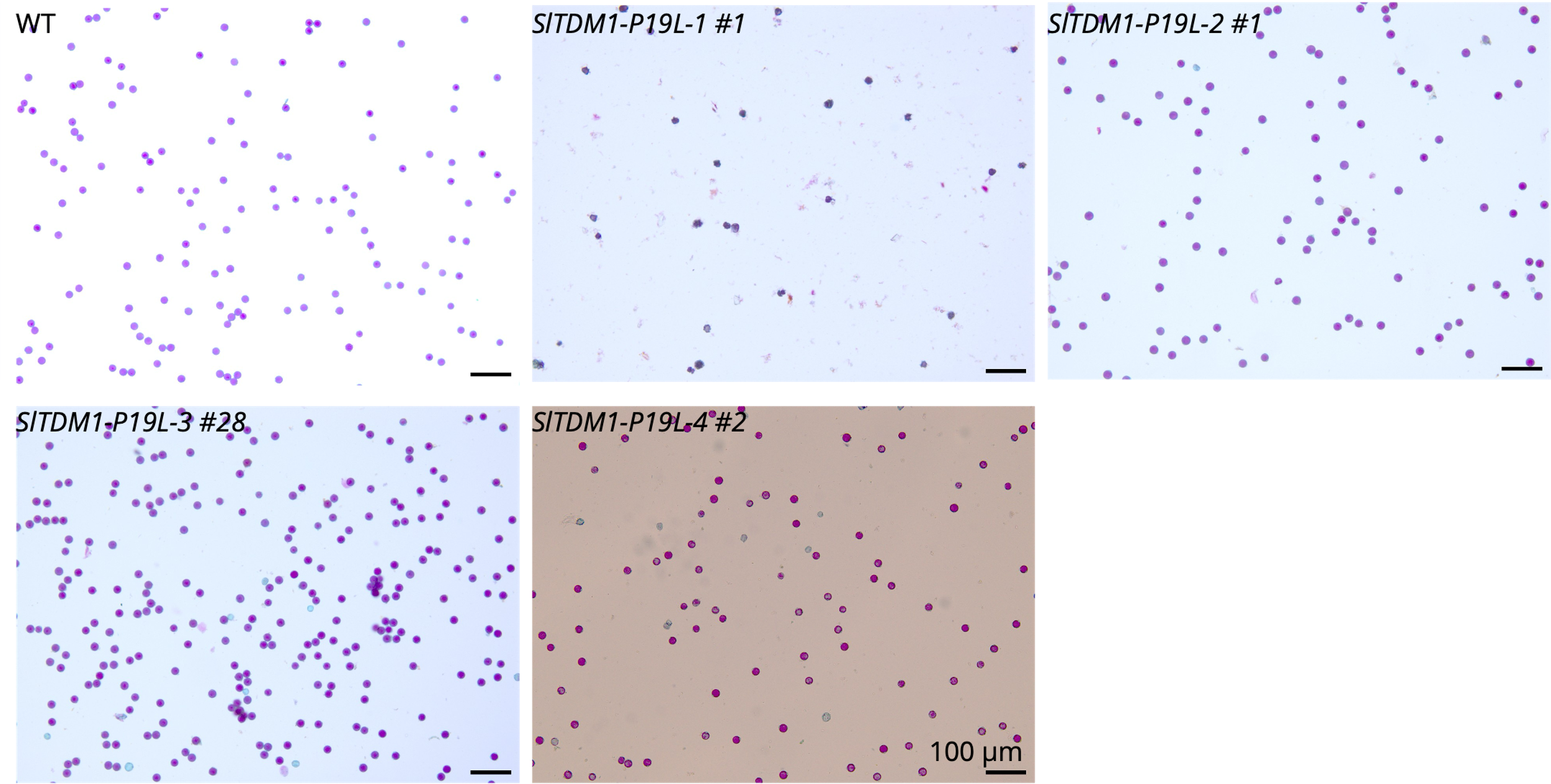

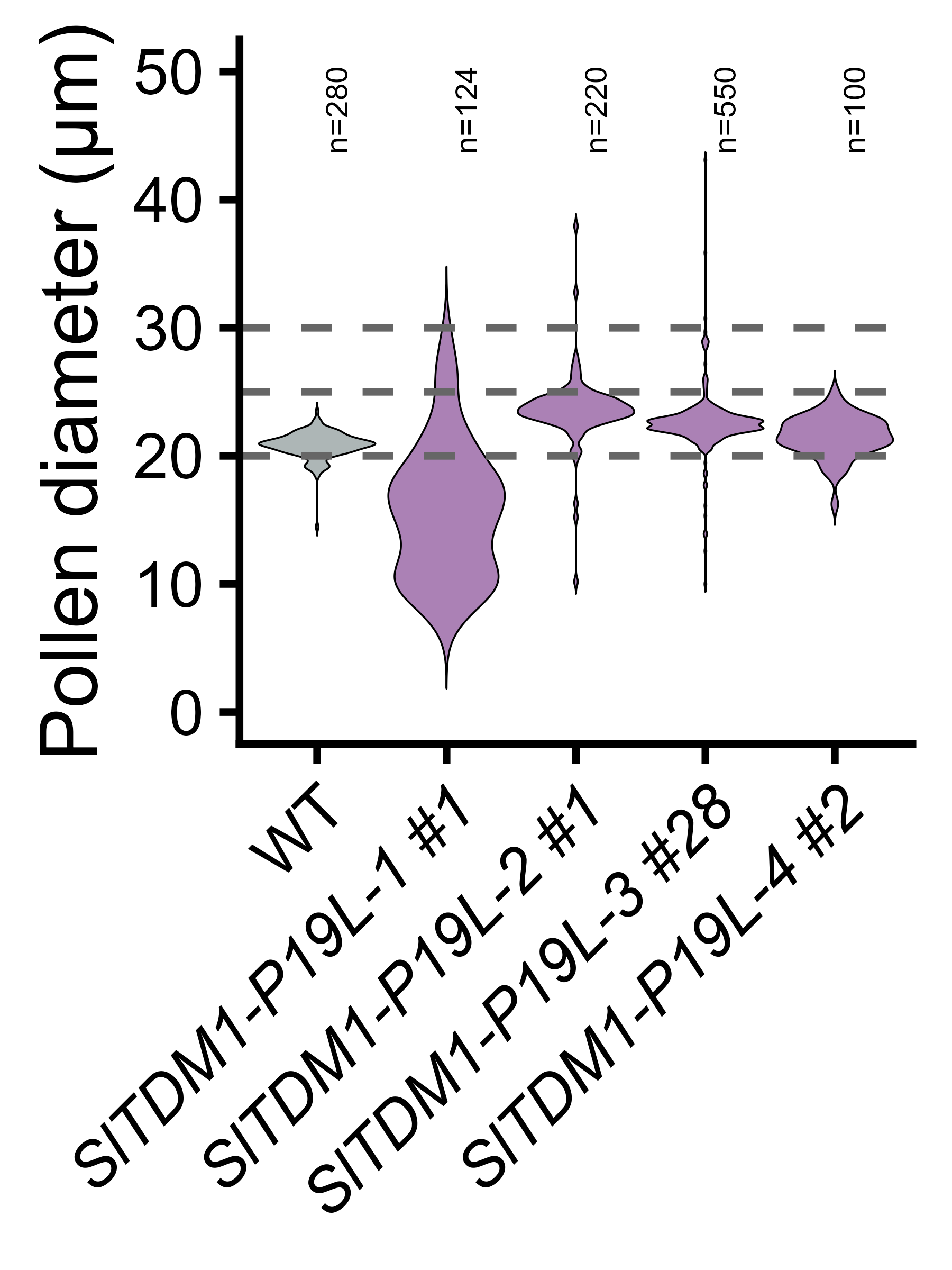

**Figure S4: Alexander staining and size quantification of pollen from diploid T1plants.**

The scale bars are 100 µm. Pollen <20 µm are deemed as nonviable, 20-25 µm as haploid and 25-30 µm as diploid. Sample sizes indicate the number of individual pollen grains measured from a single flower. WT refers to Micro-Tom.

**Table S1: Primer sequences used in this study**

| **Experiment** | **Primer pair** | **Forward primer (5'-3')** | **Reverse primer (5'-3')** |
| --- | --- | --- | --- |
| Cloning | Clone pSlTDM1 into pICH41295 | gtcgaagacaaGGAGATAATCAAGAACATTGATGAG | gcagaagactaCATTGATTCTTACGCAAAACCCTT |
|  | Clone SlTDM1 into pAGM1287 | tgtgaagacaaaATGTGGAGGAATAACGAGAGAG | gcagaagaccgcgaaTCAAGGTTTAGGACTTTCAGGA |
|  | Introduce point mutation P19L SlTDM1 | TGACCCTACCGCCGAAATGG | CCATTTCGGCGGTAGGGTCA |
|  | Domesticate SlTDM1 | GACACGAGAAAACACATCCAC | GTGGATGTGTTTTCTCGTGTC |
|  | Domesticate SlTDM1 | GTTGTCTAGGTCCCCAACAG | CTGTTGGGGACCTAGACAAC |
|  | Amplify AtTDM1 from Cifuentes et al. (2016) | GACATCGGCACTTGCTTAGAG | GCGATATAGCTCCCACTGGTT |
|  | Clone AtTMD1 into pICH47742 | agtggtctcaGGAGACATCGGCACTTGCTTAGAG | taaGGTCTCtAGCGATATAGCTCCCACTGGTT |
|  | Clone 3'UTR AtTDM1 into pAGM1301 | tgagaagacatTTCGCCTATTATCATGTACAGAA | gatgaagacgcAAGCTTGGTGCCTTTCCACTTCGT |
|  | Domesticate AtTDM1 | AGAGGAGGAGGCCACCGTAT | ATACGGTGGCCTCCTCCTCT |
|  | Domesticate AtTDM1 | GCATCTGTGAAACCGAACTC | GAGTTCGGTTTCACAGATGC |
| Genotyping | NPTII-SlTDM1 | GCATCGCCTTCTATCGCCTT | GACCCCAAGCTGGATACTCA |
|  | NPTII-AtTDM1 | GCATCGCCTTCTATCGCCTT | ACGCCGAAACCAGACAGATT |
| qPCR | qPCR ACTIN Solyc11g005330 | CATTGTGCTCAGTGGTGGTTC | TCTGCTGGAAGGTGCTAAGTG |
|  | qPCR GAPDH 3' Solyc05g014470 | TGCTGATGTCTCCGTTGTCGATC | TTTGTTGCACAAGGGACCAACC |
|  | qPCR TDM1 Solyc06g075640 | TTGTTGGGGAACTTGGCATG | GCTTGTTCTTGTCCGATTCGAG |
|  | qPCR SPO11-1 Solyc07g065440 | TTTGCAAGAGGAACCGTTGC | ACACAGGCAAATGCAGCTTG |
|  | qPCR OSD1 Solyc10g080400 | TGGTCGTCAGAACATTTCGC | TTCAATGGCCCTGACAATGG |
